# Comparative and Stability Study of Glucose Concentrations Measured in both Sodium Fluoride and Serum Separator tubes

**DOI:** 10.1101/2021.01.17.427029

**Authors:** Mustapha Dibbasey, Cathy Price, Bolarinde Lawal, Solomon Umukoro

## Abstract

**Introduction:** Sodium fluoride/potassium oxalate (NaF/KOx) tubes were once regarded as the gold-standard tubes for glucose analysis. Even though their ineffectiveness in immediately inhibiting glycolysis has been reported in several studies especially in the first 1–4hours, they are still used in our clinical Biochemistry laboratory for glucose measurement. However in its absence, only SSTs are employed for glucose measurement.

**Aim:** To determine whether SSTs can replace NaF/KOx tubes for laboratory-based measurement of blood glucose and to assess the stability of glucose concentrations for 3 days period

**Methods:** During the study period (1 March to 11 April, 2015), a total of 50 paired samples collected separately in NaF/KOx tubes and SSTs from healthy adult participants in the Gambia Adults Reference Intervals Study (GARIS) project were used as the project sample size. The samples were analysed within 2hours, and at 24hours, 42hours and 72hours time-points following blood collection and separation using Vitros 350 dry chemistry analyser. The GARIS samples were treated as clinical samples.

**Results:** There was no significant difference in the mean glucose concentrations between the two tubes (Mean difference= 0.06mmol/L; P=0.38) recorded in the different time-points. Using growth trajectory and mixed effects model, the study data showed no significant change in the glucose concentrations (p=0.25) for three days period.

**Conclusions:** The study confirms that SSTs can produce similar glucose results when employed in the absence of NaF/KOx tubes. Besides, the glucose concentrations were stable in both tubes for three days when the samples were separated within two hours and refrigerated in 2-8°C.

## Introduction

Laboratory-based measurement of blood glucose concentration is the gold-standard approach for diagnosis, treatment and assessment, and in the identification of patients at risk of developing diabetes mellitus. Diabetes mellitus (DM) is a group of metabolic disorders characterised by chronic hyperglycaemia resulting from defects in insulin synthesis, insulin action, or both (WHO, 2014). DM is a global health burden and accounts for 3.6% of the annual health budget of The Gambia (Rolfe *et al*., 1992). In 2014, WHO (2015) estimated the global prevalence of DM to be 9% among adults at least 18years of age and forecasted the disease to be the 7th leading cause of death in 2030 as the global prevalence of diabetes is projected to increase from 171 million in 2000 to 366 million in 2030 (Wild *et al*., 2000).

### NaF/KOx Tubes Effectiveness

The increase prevalence of DM has made glucose one of the most commonly requested biochemical analytes worldwide. Glucose is an unstable molecule in the whole blood due to the impact of glycolysis (Gambino, 2007), a metabolic pathway which converts glucose molecule into pyruvate (Higgins, 2007). For many decades, NaF/KOx tubes were the gold-standard collection tubes of choice when accurate glucose measurement was required. The tube type containing a combination of sodium fluoride (NaF) and potassium oxalate (KOx) was primarily designed to immediately inhibit glycolysis which negatively affects final glucose concentrations. However, several recent findings have criticised the effectiveness of NaF/KOx tubes, especially in the first 1-4hours (Chan, Swaminathan and Cockram, 1989;Roux *et al*., 2004;Peake *et al*., 2013). Mikesh and Bruns (2008) indicated that complete inhibition of glycolysis by fluoride can take as long as four hours during which time glucose can decrease as much as 0.6mmol/L at room temperature. Gambino (2009) criticised the false notion that NaF/KOx tube is an effective glycolysis inhibitor in the absence of early centrifugation following blood collection. This criticism was proven by Gambino *et al*. (2009) study which reported that the mean glucose concentration reduced by 4.6% at two hours and by 7.0% at 24 hours when blood was drawn into NaF/KOx tubes, indicating a pre-analytical loss. Furthermore, other studies including Waring, Evans and Kirkpatrick, (2007), Turchiano *et al*., 2013, and Shi *et al*., 2009 have shown that early centrifugation of blood samples in lithium-heparin and SSTs produced higher mean glucose concentration than the mean glucose concentration of NaF/KOx tubes. Elleri *et al*., (2009) reported a decrease in blood glucose by 0.47mmol/L despite samples were collected in NaF/KOx tube and iced immediately. However, these studies’ findings are contrary to the study of Spencer *et al*. (2011), which described NaF preservative to have a better stabilising effect on plasma glucose when centrifugation was performed within the first 30 minutes. Based on the findings from the studies, both ADA in 2011 and WHO discourage the use of NaF/KOx tubes to control glycolysis (Sack *et al*., 2011).

### Glucose Concentrations Stability

As part of ISO15189 accreditation process, stability study of glucose concentrations was included in the study to meet ISO15189:2012 requirements (ISO15189, 2012). Chan, Swaminathan, and Cockram (1989) further indicated that glucose concentrations were stable in NaF/KOx tubes for at least three days under room temperature following plasma separation. Al-Kharusi *et al*., (2014) reported that following plasma and serum separation, glucose concentration was stable at 4°C for seven days or even longer. In the study of Cuhadar *et al. (*2012) glucose concentrations in SSTs were stable at 4° C for up to 72 hours when centrifuged within 30 minutes. Henceforth, stability of glucose concentrations in the whole blood collected in NaF/KOx tubes for 24 hours under room temperature and in the whole blood stored at 4° C were also reported (Oddoze, Lombard and Portugal, 2012).

### NaF/KOx tubes/SSTs

The practical drawback of NaF/KOx tubes is not only its ineffectiveness in immediate glycolysis blockage but specificity to only measurement of glucose concentrations in blood and cerebrospinal fluid samples. However, SSTs have shown to be suitable for laboratory analysis of a wide spectrum of clinical chemistry analytes including blood glucose. The SSTs contain a gel which separates serum from cellular components, thus abrogating glycolysis. When compared to NaF/KOx tubes, a study conducted by Turchiano *et al*., (2013) demonstrated that mean glucose concentration for SSTs where significantly higher than the mean glucose concentration for NaF/KOx tubes whilst Frank, Shubha and D’Souza (2012) reported SSTs mean glucose concentration to be 1.15% lower than NaF/KOx mean glucose concentration when analysis was performed within 10minutes after blood collection. Gambino, (2013) demonstrated that glucose concentration in 61% of the serum samples collected in SSTs was greater than the glucose concentration in the paired plasma samples collected in NaF/KOx tubes. Following the editorial support of the study conducted by Fernandez *et al. (*2013), which reported no difference in glucose concentrations in samples collected in both NaF/KOx tubes and SSTs in field condition and separated within 2hours of blood collection, Bruns (2013) questions the need for NaF/KOx tubes for blood glucose analysis. Li *et al*., (2013) and Al-Kharusi *et al*., (2014) further suggested both NaF/KOx tubes and SSTs to be suitable candidates for glucose measurement. Currently, SSTs are used for laboratory-based measurement of blood glucose in the absence of NaF/KOx tubes in our Biochemistry laboratory. As such, the study aims to determine whether glucose concentrations will be varied significantly in the paired NaF/KOx and SSTs samples and to assess the stability of glucose concentrations for 72hours.

### Aim

To determine whether SSTs can replace NaF/KOx tubes for laboratory-based measurement of blood glucose and to assess the stability of glucose concentrations for 3 days period

### Hypothesis

H_0_= there will be no significant difference in the glucose concentration measured using both NaF/KOx tubes and SSTs

H_1_: there will be a significant difference in the glucose concentration measured using both NaF/KOx tubes and SSTs

## Materials and Methods

### Sample Collection/Handling

The study was conducted in the Medical Research Council’s (MRC) Biochemistry laboratory. With the permission of the study investigators, the blood samples for the practical project were obtained from the GARIS project which started in March 2015, an on-going project targeting to establish haematological and biochemical reference values for Gambian adults in Greater Banjul Area. During the practical project (1 March to 11 April, 2015), a total of 50 pairs samples collected separately in NaF/KOx and SSTs from healthy adult participants in the GARIS project were used as the project sample size.

The participants who were healthy adult donors (≥18 and ≤45years) residing in the urban areas and negative for HIV, VDRL and Hepatitis B were enrolled into the GARIS project. The participants were bled and the blood samples were received, in the clinical laboratories under room temperature.

### Inclusion criteria

The first 50 pairs of GARIS samples with sufficient volume were used in the practical project.

### Sample analysis

Prior to the analysis, quality controls were run to ascertain the functionality of the Vitros 350 Analyser, which performed the glucose analysis of the project samples. Centrifugation (separation of plasma from NaF blood and serum from clotted blood samples) and first analysis of glucose concentrations were performed within two hours of blood collection, and then the serum and plasma samples were stored at 2-8° C temperature. The subsequent analyses were exactly performed at the following time-points: 24, 48 and 72 hours. The study samples were analysed using the same lot of reagent, eliminating any lot-to-lot variability in the results.

### Statistical analysis

The statistical analysis was performed using STATA version 12 and Microsoft Word excel. Paired t-test was performed to determine any significant statistical difference in the mean glucose concentration recorded in the time-points. Pearson correlation was performed to determine the association between NaF/KOx tubes and SSTs glucose concentrations. A mixed effects model and growth trajectory were performed to assess the stability of glucose concentrations.

### Ethical Issues

The GARIS project was approved by MRC Scientific Coordinating Committee and the Gambia Government/MRC Joint Ethics Committee (MRC, 2013) prior to commencement of the study. Written informed consent was obtained from study participants enrolled in the GARIS study. GARIS participants were assigned unique identification numbers for maintaining anonymity and confidentiality.

## Result

Table:1 and figure:1 shows mean glucose concentrations (MGC) measured in both NaF/KOx tubes and SSTs at different time-points. The total mean concentrations of NaF/KOx tubes (6.88±4.36 mmol/L) and SSTs (6.94±4.31 mmol/L) produced an overall difference of 0.06mmol/L [Total Mean difference (NaF/KOx-SST) = 0.06±0.07; n=50]. When the MGC from the time-points were compared using two-tailed paired t-test, the result shows no significant difference (P=0.38). The Pearson correlation (Figure: 2) shows high significant correlation (R=0.9999 P<0.00001) between NaF/KOx tubes and SSTs glucose concentrations. The P value less that 0.05 indicates a statistical significant difference.

**Table 1.** Showing MGC of NaF/KOx tubes and SSTs from different time-points

| Time-points | NaF/KOx MGC | SST MGC | MD (NaF/KOx-SST) |
| --- | --- | --- | --- |
| within 2hrs | 6.86 ±4.31 | 6.92 ±4.29 | -0.06 |
| 24hrs | 6.88 ±4.34 | 7.01 ±4.35 | -0.13 |
| 48hrs | 6.87 ±4.38 | 7.01 ±4.42 | -0.14 |
| 72hrs | 6.92 ±4.38 | 6.82 ±4.30 | 0.10 |
| **Total mean** | **6.88±0.07** | **6.94 ±0.07** | **-0.06** |

MGC: mean glucose concentration, MD: mean difference (i.e. differences between the time-points means), SST: serum separator tubes, NaF/KOx: sodium fluoride tubes

**Figure 1.**
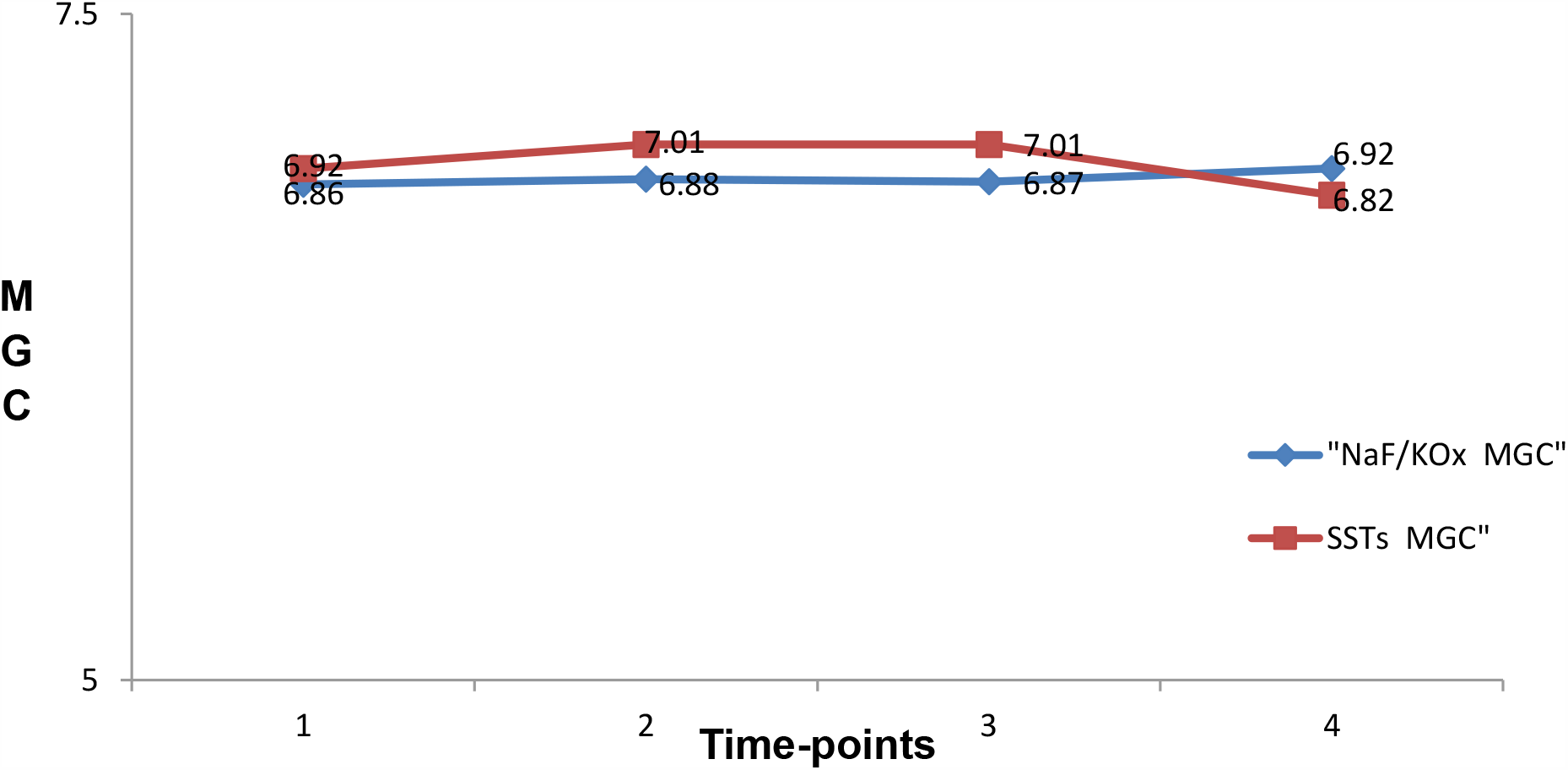
Graph showing the MGC of NaF/KOx tubes and SSTs recorded in the time-points 1 (2hours), 2 (24hours), 3 (48hours) and 4 (72 hours)

**Figure 2.**
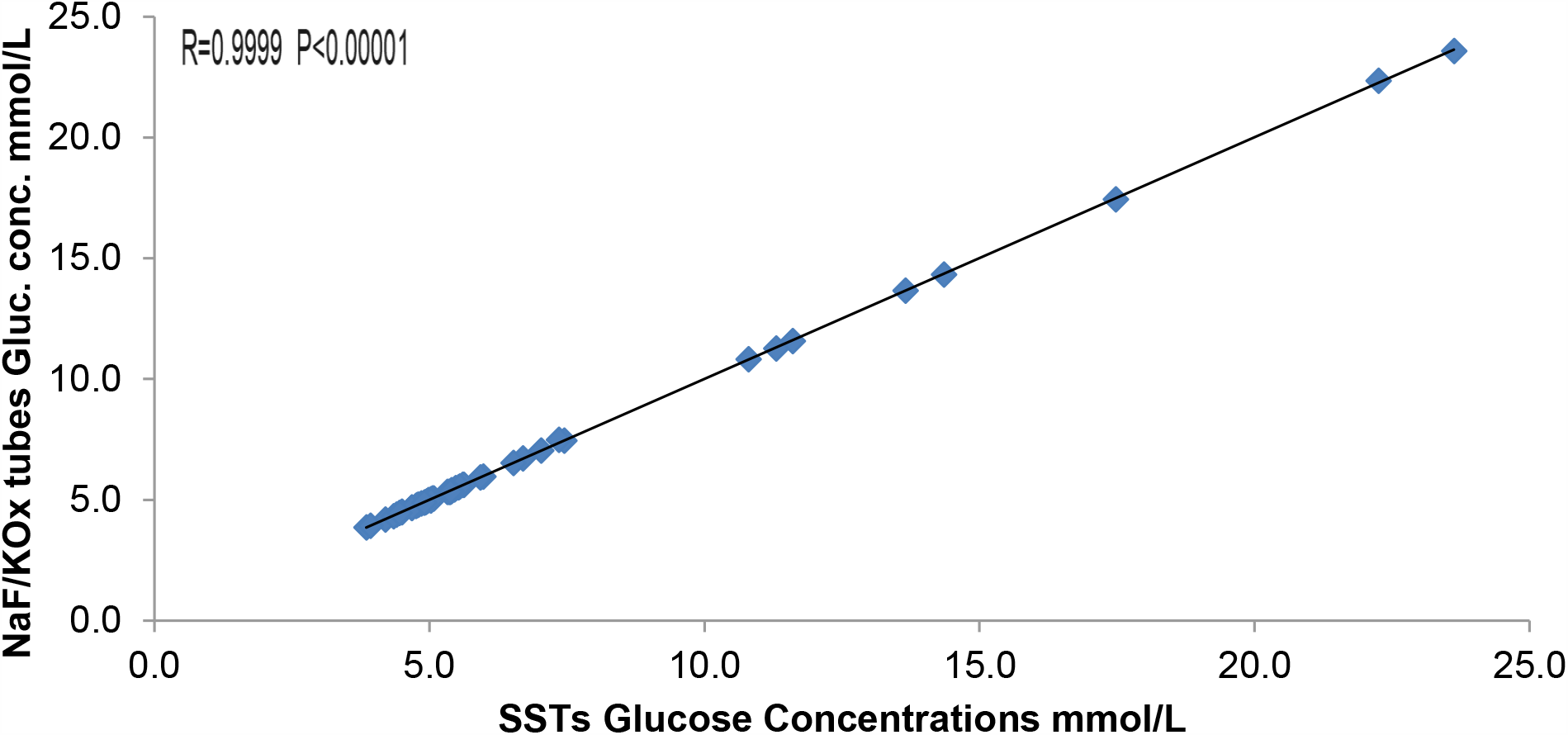
Graph showing relationship between glucose concentrations meaured in NaF/KOx tubes and SSTs

Growth trajectory graph (Figure:3) was used to determine the stability of individual glucose concentrations in both tubes after the first analysis and up to 72 hours of analysis. The graph shows that the glucose concentrations measured with NaF/KOx and SSTs were stable from the first analysis up to 72 hours at 2-8° C.

**Figure 3.**
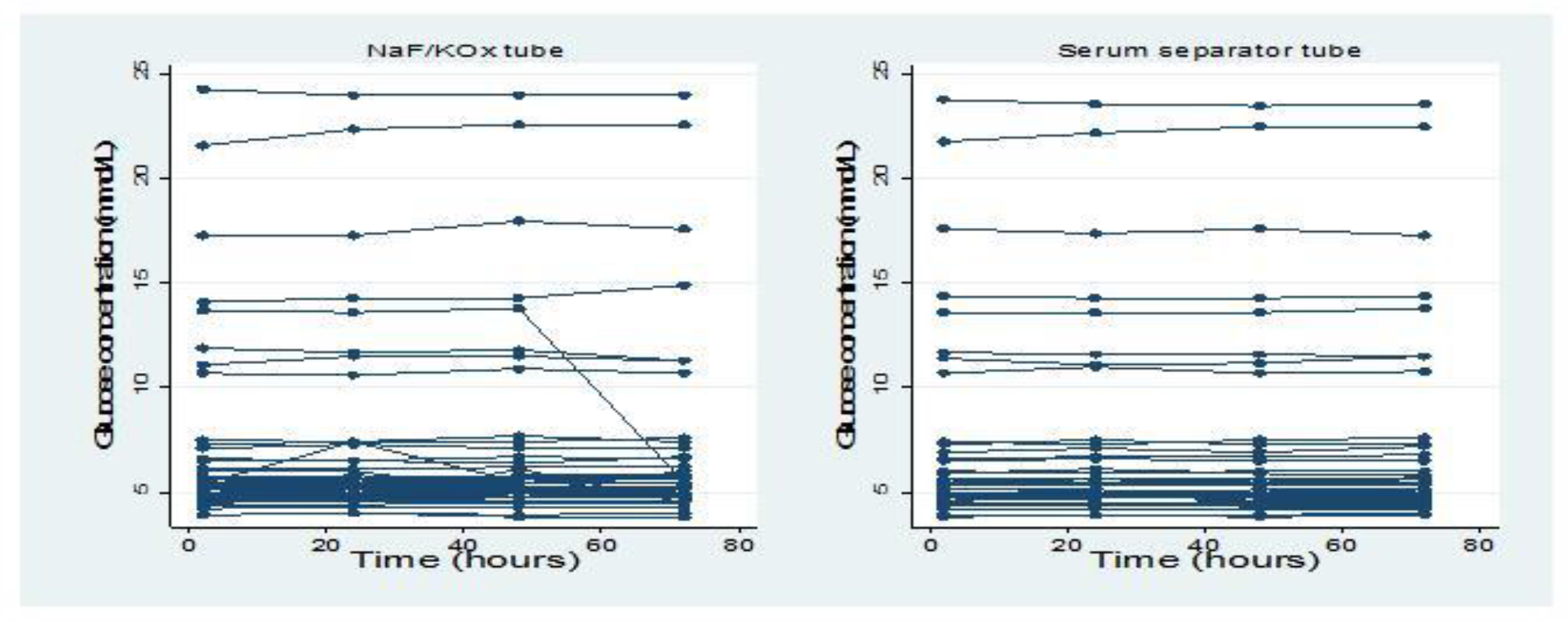
Growth trajectory graph for determining the stability of individual glucose concentrations from 2 to 72 hours

For statistical analysis purpose, raw glucose concentrations were transformed using the inverse of the square of the glucose concentration. Post-data transformation, a mixed effects model (Table:2) was employed for stability study of glucose concentrations in both tubes using within 2hours as the reference category. There was no evidence of change in the glucose concentration (overall p=0.25) in both tubes for 3 days.

**Table 2.** Stability study of glucose concentrations from the 2 hours to 72 hours of analysis

| Time-points | Estimated difference (95% CI*) | p. value |
| --- | --- | --- |
| Within 2 hours" | 0 |  |
| 24 hours | 0.001 | 0.06 |
| 48 hours | 0.0003 | 0.33 |
| 72 hours | 0.0002 | 0.67 |
| **Time** |  | **0.25#** |

\*: 95% confidence interval.
#: Overall P. value

## Discussion

The study demonstrates no significant difference in the mean glucose concentrations between NaF/KOx tubes and SSTs. The study outcome is consistent with the outcome of the studies of Fernandez *et al. (*2013), Li *et al*., (2013) and Al-Kharusi *et al*., (2014), which qualify both tubes to be suitable candidates for glucose analysis. The slight increase in the SSTs MGC (0.06mmol/L) as compared to NaF/KOx tubes’ explains the ineffectiveness of NaF/KOx tubes and that the use of NaF/KOx tubes may impact DM diagnosis especially for patients in borderline area of impaired glucose tolerance. Furthermore, this study, and other comparative and review studies have proven that the gold-standard tube is ineffective in its role to instantaneously block glycolysis (Gambino, 2007;Gambino *et al*., 2009;Mikesh and Bruns, 2008;Gambino, 2013;Bruns, 2013;Gambino and Bruns, 2013), hence its usage is being discouraged by ADA (Sack *et al*., 2011).

Since both tubes are suitable candidates as shown by the data (Table:1&Figure:1), preference is given to SSTs as it can be used to analyse wide spectrum of biochemical analytes, and avoid inconveniencies and reduce mistakes associated with analysing both plasma and serum at the Biochemistry laboratory for an individual patient. Clinically, the use of only SSTs will reduce the volume of blood collected from patients when glucose and other analytes are requested together and improve glucose turn-around-time as well as laboratory workflow. Even with the low cost, elimination of NaF/KOx tubes would be a significant cost-saving measure for the National Health Service as large number of laboratory-based glucose tests are performed annually.

However, this study result differs from the previous studies including Gambino (2013), Waring, Evans and Kirkpatrick, (2006), Turchiano *et al*., (2013) and Shi *et al*., (2009), which reported higher significant differences in the MGC of SSTs as compared to the MGC of NaF/KOx tubes. The significant difference was due to early separation as SSTs blood samples were centrifuged immediately after collection whereas NaF/KOx samples were centrifuged immediately before glucose analysis. Since blood cells (erythrocytes, leucocytes and thrombocytes) are known to metabolise blood glucose, Mikesh and Bruns (2008) indicated that early separation of plasma/serum from the blood cells instantaneously abrogates glycolysis as the blood cells are prevented from metabolising blood glucose to glucose-6-phosphate and other phosphates. However, Frank, Shubha and D’Souza, (2012) found SSTs MGC to be lower than NaF/KOx MGC when analysis was performed within 10 minutes after blood collection.

### Glucose Concentrations Stability

After the first analysis, glucose concentrations were stable at 2-8° C in both tubes for three days. Similar finding was reported by Chan, Swaminathan and Cockram, (1989) even though the samples were stored in room temperature, Cuhadar *et al. (*2012) and Al-Kharusi *et al*., (2014). The stability of glucose concentrations after four hours in NaF/KOx tubes was explained in the study of Mikesh and Bruns (2008) which reported that NaF inhibits enolase within five minutes of addition to the blood while enzymes in the upstream of glycolysis pathway remain active, allowing continuous metabolism of glucose by blood cells into glucose-6-phosphate and other phosphorylated metabolites until there is no available supply of ATP in the cells. Supply of ATP is exhausted about 60 minutes after addition of NaF and then glucose concentration becomes stabilised in the tube (Gambino, 2007). In the case of SSTs, Bruns (2013) explains that SSTs contains gel barrier that separates blood cells from plasma, thus this means that glycolysis is completely ceased as soon as separation is performed. It further explains the stability of glucose concentrations in the SSTs (Shi *et al*., 2009;Mikesh and Bruns, 2008). The stability study hence proves that glucose concentration can be measured in plasma/serum within 3days when the samples are refrigeration at 2-8°C.

### Limitations

The study failed to address potential confounders such as diabetes status that can influence the observed results. From the study of Spencer *et al*. (2011), the data showed that plasma glucose in NaF/KOx reduced by 16.4 in non-diabetic and by 10.4 in diabetic patients.

Furthermore, it was not possible to estimate the effects of red blood cells (RBCs) haemolysis on glucose concentrations compared between the two tubes in the study. The studies of Fernandez *et al*., (2013), Al-Kharusi *et al*., (2014) and Ko *et al*., (2015) all reported high RBCs haemolysis in NaF/KOx tubes than in SSTs samples but failed to assess the impact of the haemolysis. This high haemolysis rate observed can potentially influence glucose concentration and might have resulted in the slight decrease in NaF/KOx mean glucose concentration by 0.06mmol/L. Catalase released from the lysed erythrocytes can negatively affect glucose values (Lippi *et al*., 2006).

## Conclusion

The study confirms that SSTs can be satisfactorily employed in the place of NaF/KOx tubes for laboratory glucose measurement in our settings. Accepting the use of SSTs for glucose measurement offers numerous operational benefits including cost-saving measure for the National Health Service. The glucose concentrations were stable in both tubes for three days when the plasma/serum samples were refrigerated at 2-8_°_C. Further study is required to provide an insight into the effects of haemolysis on the glucose concentrations and its stability whilst considering potential confounders.

## Abbreviations

SSTs: Serum Separator Tubes
NaF/KOx: Sodium fluoride/potassium oxalate
WHO: World Health Organisation
ADA: American Diabetes Association
EDTA: Ethylenediamine tetra acetic acid
GARIS: Gambia Adults Reference Interval Study
NaF: Sodium Fluoride
MRC: Medical Research Council
ATP: Adenosine Triphosphate
GARIS: Gambia Adult Reference Interval Study
HIV: Human Immunodeficiency Virus
VDRL: Venereal Disease Research Laboratory
RBCs: Red Blood Cells
≤: Less than or equal to
≥: Greater than or equal to

## Notes

### Competing Interest Statement

The authors have declared no competing interest.

